# Plantarflexor Moment Arms Estimated From Tendon Excursion In Vivo Are Not Well Correlated With Geometric Measurements

**DOI:** 10.1101/290916

**Authors:** Josh R Baxter, Stephen J Piazza

**Author notes:** Mailing address.* 3737 Market Street, Suite 702, Philadelphia, PA, USA 19104. Corresponding author.* Josh R. Baxter, PhD.

## Abstract

Geometric and tendon excursion methods have both been used extensively for estimating plantarflexor muscle moment arm *in vivo.* Geometric measures often utilize magnetic resonance imaging, which can be costly and impractical for many investigations. Estimating moment arm from tendon excursion measured with ultrasonography may provide a cost-effective alternative to geometric measures of moment arm, but how well such measures represent geometry-based moment arms remains in question. The purpose of this study was to determine whether moment arms from tendon excursion can serve as a surrogate for moment arms measured geometrically. Magnetic resonance and ultrasound imaging were performed on 19 young male subjects to quantify plantarflexor moment arm based on geometric and tendon excursion paradigms, respectively. These measurements were only moderately correlated (R^2^ = 0. 21, *p* = 0.052), and moment arm from tendon excursion under-approximated geometric moment arm by nearly 40% (p < 0.001). This moderate correlation between methods is at odds with a prior report (N = 9) of a very strong correlation (R^2^ = 0.94) in a similar study. Therefore, we performed 92,378 regression analyses (19 choose 9) to determine if such a strong correlation existed in our study population. We found that certain sub-populations of the current study generated similarly strong coefficients of determination (R^2^ = 0.92), but 84% of all analyses revealed no correlation (p > 0.05). Our results suggest that the moment arms from musculoskeletal geometry cannot be otherwise obtained by simply scaling moment arms estimated from tendon excursion.

**Funding:** No funding was provided for this research

**Conflict of interest:** The Authors have no conflicts of interest related to this study to report

**Acknowledgements:** The Authors would like to thank the Social, Life, and Engineering Sciences Imaging Center at Penn State University for providing imaging time.

## Introduction

Plantarflexor moment arm is an important musculoskeletal parameter that simultaneously determines muscle mechanical advantage and influences the amount of muscle fiber shortening or lengthening that occurs during a given joint rotation. *In vivo* measurements of plantarflexor moment arm are often made using one of two methodological paradigms: 1) geometric methods that quantify the distance between the tendon line of action and the joint axis of rotation [1]; and 2) tendon excursion methods based on the Principle of Virtual Work that consider plantarflexor moment arm to be equal to the amount of tendon travel occurring per unit of joint rotation [2]. While many investigators have made use of magnetic resonance (MR) imaging to perform geometric measurements of muscle moment arm [1,3,4], high cost and limited access to this imaging modality are important barriers to its implementation in some laboratories. Conversely, moment arm estimation using tendon excursion measured with ultrasonography (US) is a less expensive and more easily implemented approach [5,6]. However, muscle-tendon compliance introduces uncertainty to measurements that results in under approximations of moment arm when compared to geometric measurements made using MR imaging [5].

Despite many reports of plantarflexor moment arms in the literature using either geometric or tendon excursion methods, direct comparisons between moment arms measured with geometric/MR and tendon excursion/US measurements in the same individuals have been reported in only a single study [5]. Fath and colleagues found that moment arms measured with tendon excursion/US strongly correlated (R^2^ = 0.94) with those measured in the same nine participants using geometric/MR measures. While these initial findings support the possibility that tendon excursion/US is a good surrogate measure of geometric/MR, this strong correlation has yet to be replicated and may potentially have resulted because of small sample size. Tendon excursion/US measures were also found to underestimate by nearly 30% those measured with geometric/MR measures; which may be an artifact of tendon relaxation caused by variations in tendon slack length or compliance. Regardless of these differences, the finding of such a strong correlation suggests that tendon excursion/US may be a viable surrogate for geometric/MR measures of plantarflexor moment arm. Before reaching this conclusion, however, it is important to see this finding replicated in a larger number of participants.

The purpose of this study was to compare plantarflexor moment arm measurements when both geometric/MR and tendon excursion/US methods were applied in the same participants. Based on the previous study of Fath et al. [5], we expected to find that tendon excursion/US moment arms are viable stand-ins for geometric/MR measures of moment arms. Specifically, we hypothesized that tendon excursion/US measures of moment arm will be strongly correlated with geometric/MR measures, but that tendon excursion/US moment arms will be systematically smaller. Further support for this hypothesis could lead to tendon excursion/US becoming a low-cost and reliable alternative to the geometric/MR method with the limitation that moment arm measurements would be correlated but under approximated compared to geometric/MR.

## Methods

Plantarflexor moment arm was quantified using geometric/MR and tendon excursion/US measurements in 20 healthy young males (age: 26.0 ± 3.5 y, stature: 177.7 ± 7.7 cm, and body mass: 76.3 ± 15.6 kg). MR images of the lower leg and foot were acquired in a 3.0 T MR scanner [3], and tendon excursion was measured using US [5]. Measurement techniques for both geometric/MR and tendon excursion/US were similar to those employed by Fath and colleagues (2010) in order to make direct comparisons. All procedures involving human participants were approved by the Institutional Review Board of The Pennsylvania State University.

Magnetic resonance images of the right lower leg and foot were acquired with the foot passively resting in neutral position (0°), 10° plantarflexion, and 10° dorsiflexion using a 3.0 T scanner (Siemens; Erlangen, Germany) and established scanning parameters (three-dimensional isotropic T1 weighted sequence; echo time: 1.31 ms, repetition time:3.96 ms, 3.96 mm, 500-mm field of view, 0.9 mm voxel size; [7]. Subjects rest on the scanning bed in the supine position with both knees fully extended. Images of a quasi-sagittal plane containing the midline of the Achilles tendon and the longitudinal axis of the second metatarsal were reconstructed by a single investigator using an open-source medical imaging viewer (Osirix, Pixmeo, Geneva, Switzerland) and printed onto transparent sheets. The instantaneous center of rotation between the tibia and talus in the sagittal plane at neutral position was calculated using Reuleaux’s method [8] from 10° dorsiflexion to 10° plantarflexion [7]. The Achilles tendon line of action was defined as the midline of the free tendon in the image acquired in the neutral position. Plantarflexor moment arm was calculated as the shortest distance between the center of rotation and tendon line of action in the neutral position.

Tendon excursion measurements were performed using US while ankle rotations were controlled using a dynamometer [5] while subjects were seated in the device with the knee fully extended. Briefly, the medial gastrocnemius muscle-tendon junction of the right leg was tracked under B-mode ultrasonography (Aloka 1100; transducer: SSD-625, 7.5 MHz and 39-mm scan width; Wallingford, CT) as the foot was moved from 10° dorsiflexion to 30° plantarflexion at a rate of 10°s^−1^. Ultrasound images were acquired and saved to a personal computer at 30 Hz using a frame grabber card (Scion Corporation, LG-3, Frederick, MD, USA). The location of the muscle-tendon junction was manually digitized on US images, and the plantarflexor moment arm was defined as the slope of a best-fit line of the muscle tendon position as a function of ankle angle between 10° dorsiflexion and 10° plantarflexion (Figure 1), which was found to generate the strongest correlation with geometric/MR measurements in a previous study [5]. The probe was aligned on the leg to ensure that the muscle-tendon junction was clearly visible throughout ankle rotation. Once aligned, the US probe was secured to the posterior aspect of the lower leg using a foam cast and self-adhesive wrap. A wire taped to the skin near the muscle-tendon junction cast a shadow that was used to confirm that the probe did not move with respect to the skin during foot rotation. Images acquired using the frame grabber were found to be corrupted in one subject; thus, comparisons between tendon excursion/US and geometric/MR were performed in the remaining 19 subjects.

**Figure 1.**
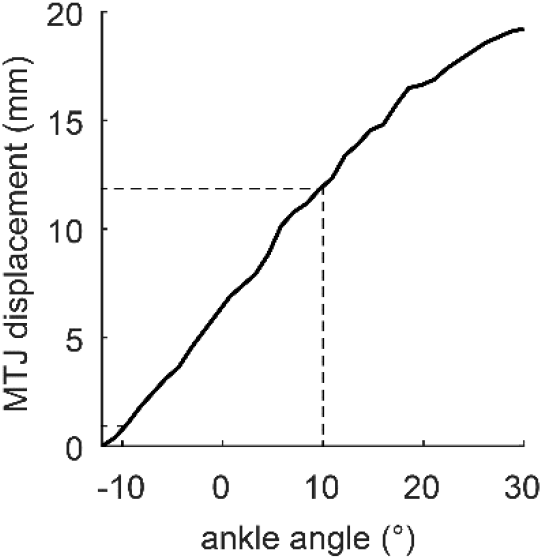
Muscle-tendon junction (MTJ) displacements were manually digitized and plotted against ankle angle to quantify tendon excursion/US. This representative image showed a strong correlation (R^2^ = 0.997) between MTJ and ankle angle between 10° dorsiflexion (-) and 10° plantarflexion (+).

Measurements of plantarflexor moment arm from geometric/MR and tendon excursion/US were correlated using simple linear regression. Moment arm magnitudes were compared between techniques using a paired t-test (α = 0.05). A Bland-Altman plot [9] was constructed to visually assess the correspondence between the two measurements. We considered the geometric/MR measurements to be a ‘gold standard’ because it clearly defines both the tendon line of action and ankle center of rotation. After this initial assessment, we performed 92,378 linear correlations (19 choose 9) in an attempt to recreate the very strong correlation between tendon excursion/US and geometric/MR methods using a smaller sample size previously reported [5].

## Results

Plantarflexor moment arm estimated using tendon excursion/US was only moderately correlated with tendon-excursion measurements explaining 21% of the variance in geometric/MR moment arm measurements that approached statistical significance (Figure 2A, R^2^ = 0.21; p = 0.052). Paired t-tests revealed that tendon excursion significantly under-approximated moment arms calculated using geometric methods by nearly 40% (Table 1, *p* < 0.001). A Bland-Altman plot similarly illustrated this under-approximation and demonstrated that this error tended to be larger for those subjects who have smaller moment arms (Figure 3, R^2^ = 0.17, p = 0.079).

**Figure 2.**
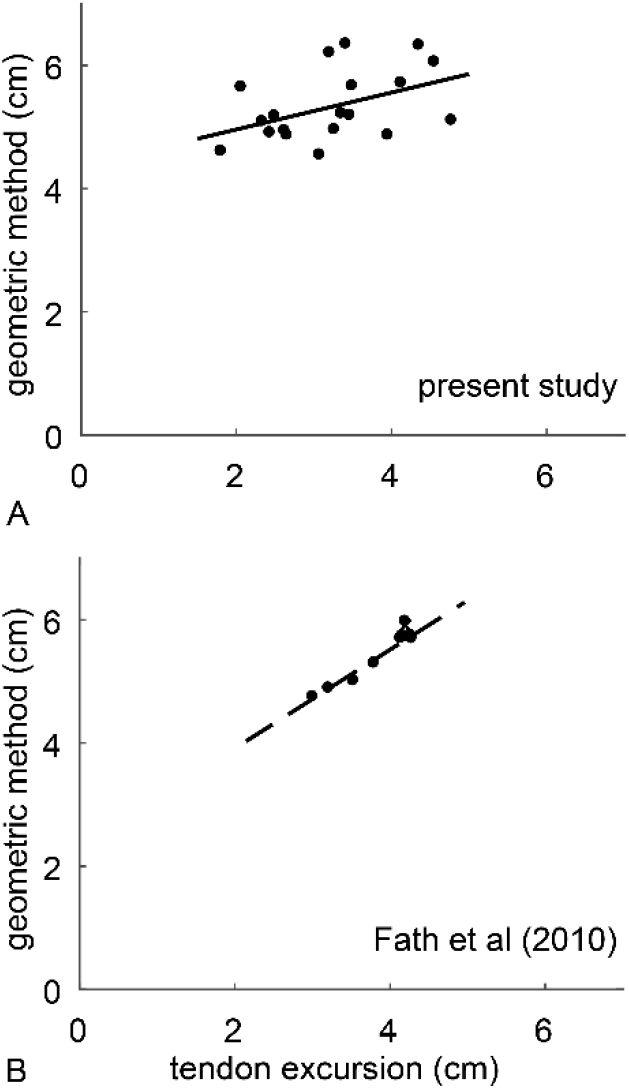
(A) Plantarflexor moment arms in the present study measured using geometric and tendon excursion methods in the same subjects showed a moderate-positive correlation that approached statistical significance (R^2^ = 0.21; p = 0.052). (B) A prior report by Fath et al. (2010) found a very strong positive correlation (R^2^ = 0.94) between the measures of moment arm.

**Figure 3.**
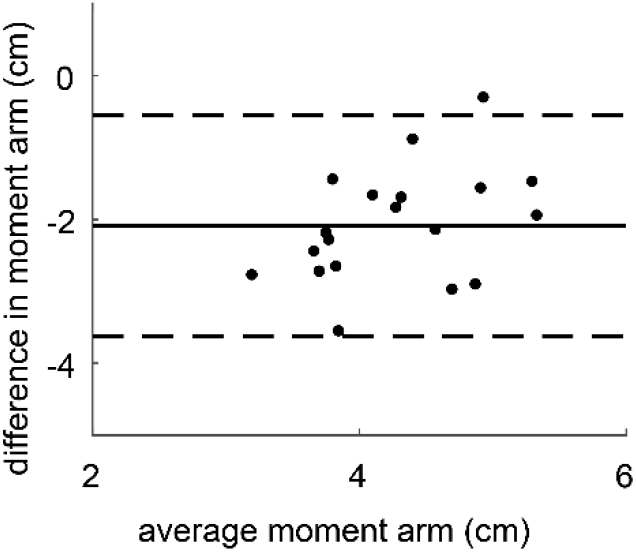
Bland-Altman plot showing the systematic under-approximation of moment arm when measured by tendon excursion. If tendon excursion were a surrogate measure of geometric methodology, then the variability in measurement differences (dashed lines are 1.96 x standard deviation) would be small and centered around zero (mean differences given by solid line).

**Table 1.** Mean (with 95% confidence interval) values for plantarflexor moment arm of the Achilles tendon calculated using geometric and tendon excursion methods. *Difference between methods was statistically significant (p < 0.001). Values from the present study and that of Fath et al. (2010).

|  | Geometric Method | Tendon Excursion | % difference |
| --- | --- | --- | --- |
| Present study | 5.3 cm (5.1 – 5.6 cm) | 3.2 cm (2.9 – 3.6 cm) | -40%* |
| Fath et al. | 5.4 cm (5.1 – 5.7 cm) | 3.8 cm (3.5 – 4.2 cm) | -30%* |

Regression analyses on sub-populations (N = 9) of the overall study-population (N = 19) yielded a maximum coefficient of determination (R^2^) of 0.92. Of the 92,378 regression analyses (Figure 4), only 11% yielded strong coefficients of determination (R^2^ > 0.50), only 16% were statistically significant correlations (p < 0.05), and nearly half of all sub-populations explained less than 20% of the variance in moment arm (R^2^ < 0.20).

**Figure 4.**
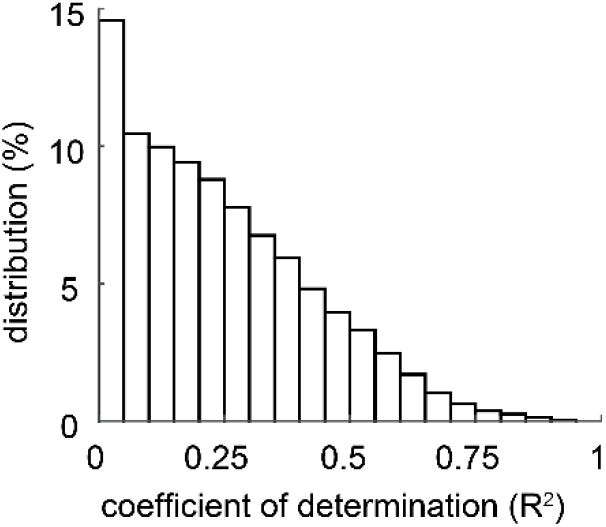
Histogram showing the coefficient of determination (R^2^) distribution when subsamples (N = 9) were analyzed to compare with literature values. Linear regressions performed on 92,378 sub-populations of the 19 subjects (19 choose 9) show that 11% of combinations resulted in strong correlations (R^2^ > 0.50).

## Discussion

The results of the present study do not support our hypothesis that plantarflexor moment arms measured *in vivo* using tendon excursion/US would be strongly correlated with those measured using the geometric/MR method. While we did find a moderate correlation between the two measures that approached statistical significance (R^2^ = 0.21, p = 0.052), our findings suggest that tendon excursion/US explained only a small fraction of the variance in geometric/MR moment arms, suggesting that tendon excursion/US were affected by measurement artifact that may have resulted from inter-subject differences in tendon slack length and compliance.

The plantarflexor moment arms we measured using geometric/MR and tendon excursion/US methods compared favorably to values previously reported (Table 1). MR imaging studies have yielded moment arm values for young adults ranging from 4.8 to 6.0 cm [1,3–5,10], which closely matched our 95% confidence-interval of 5.1 to 5.6 cm. Our geometric measurements also agreed with dynamic measures of moment arm that were acquired during gait using a combined ultrasound and motion capture approach [11]. Previously reported tendon excursion/US measurements of plantarflexor moment arm in young men range from 3.1 to 4.2 cm [5,12–14], similar to our 95% confidence-interval values of 2.9 to 3.6 cm.

Our primary finding that plantarflexor moment arm measured using tendon excursion/US is only moderately correlated with geometric-based measurements is at odds with the previous results of Fath et al. (2010) (Figure 2). While our sample size (N = 19) was larger than a prior report by Fath et al. (N = 9), our data were not adequately powered (β = 0.55), suggesting that the relationship between tendon excursion/US and geometric/MR is not as strong as previously reported. Increasing our sample size further would have increased the statistical power but may not have changed the coefficient of determination. A Bland-Altman plot (Figure 3) also demonstrates that the disagreement between measurements increases with measurement magnitude. Both study populations were healthy young adults with moment arm measurements that were similar across studies for each measurement method (Table 1). MR and US images were acquired and processed using similar methodology to that employed by Fath et al. (2010). To the best of our understanding, the primary difference between these two studies was the sample size, with the current study having more than twice as many participants. When we performed regressions using all combinations of our data set to match the sample size of the prior study, we found rare instances with similar coefficients of determination (Figure 4).

Inter-individual variation in tendon compliance and slack length also has the potential to affect tendon excursion during *in vivo* measurements. Passive loading of the Achilles tendon occurs even in plantarflexion and increases with ankle dorsiflexion [5], which violates a key assumption of the Principle of Virtual Work [2] that requires tendon energy storage (and therefore tension) to remain constant throughout the measurement. Tendon compliance and slack length dictate tendon elongation (and therefore tension) during passive ankle movements [15] and demonstrate variability within healthy adult cohorts [16,17]. Directly tracking the calcaneal insertion of the Achilles tendon with MR produces moment arm measurements similar to geometric/MR [4], suggesting that variations in tendon parameters explain differences between tendon excursion/US and geometric/MR. Other groups have attempted to control for Achilles tendon loading by having subjects actively contract the plantarflexors during the measurement [13,14]. We similarly attempted to control for tendon loading by measuring tendon excursion with US as subjects either maintained a constant 30%, 60%, or 100% of their maximal isometric torque. We found that these attempts to control for tendon tension resulted in moment arm approximations that did not significantly correlate with geometric/MR measurements (all R^2^ < 0.09, p > 0.21, Appendix Table 1), but it is possible that a different means of controlling tendon tension in vivo might result in stronger correlations between moment arms measured using the two methods. Further, we correlated other geometric-based moment arm measurements [18–20] with geometric/MR and found that these techniques compared more favorably than tendon excursion/US (Appendix Table 1).

This study has certain limitations that should be considered in the interpretation of our findings. Ankle rotation was not tracked with motion capture, instead we visually confirmed that the foot remained flat on the dynamometer foot plate throughout testing. Our assessments of moment arm using the geometric/MR technique were limited to neutral position because our method required location of the center of rotation from images made in the 10° plantarflexed and dorsiflexed positions. It is possible that the correlation between the two measures depends on joint angle as geometric/MR moment arm tends to increase with plantarflexion [4], but we did not test this possibility. We quantified the geometric/MR moment arm in a quasi-sagittal plane rather than analyzing the three-dimensional moment arm that may provide a more accurate measurement [21,22]. Finally, our study population was limited to healthy young males, a choice we made to minimize the possible effects of variation due to age and sex on moment arm, but one that limits our conclusions to this specific population.

Based on our findings, we recommend that measurements of moment arm based on geometry be used when possible. Geometric measurements rely on fewer assumptions than measurements based on tendon excursion, but when geometry is assessed with MR this approach can be costly and impractical. Advances in MR imaging provide the capability to quantify the three-dimensional moment arm of the Achilles tendon [21,22], which may have implications on pathologic populations [23]. As an alternative, three-dimensional motion capture and US have been combined to permit assessment of the distance between the transmalleolar axis and the Achilles tendon line of action during functional activities [11,24]. Quantifying moment arm dynamically in this way may provide new insight into the implications of joint structure [3,25] and muscle activity [11] on joint leverage and function.

## Appendix

### Correlating other measurements of moment arm with geometric/MR

A secondary aim of this study was to correlate plantarflexor moment arms assessed using various other reported techniques or surrogate measures of with geometric/MR measurements measured in the same subjects (Appendix Table 1). External measurements of the malleoli were acquired using digital photography (Scholz et al., 2008). When tendon tension was controlled in tendon excursion experiments, this was done by instructing the subjects to generate a desired amount of ankle torque during each ankle rotation. This feedback was provided by a computer display showing a dial with both the desired torque and actual torque. Subjects were provided practice trials until they were able to steadily maintain the desired torque. These tendon excursion/US measurements were acquired after subjects performed three maximal voluntary isometric contractions (MVIC), which were used to generate subject specific torque targets of 30%, 60%, and 100% MVIC.

Geometric measurements of moment arm all positively correlated with geometric/MR. The geometric/MR measurement of the instantaneous center of rotation between the tibia and calcaneus was strongly correlated (R^2^ = 0.90) and had an RMSE less than 2 mm. This analysis was performed to confirm that the geometric/MR measurement works similarly well when tracking either the talus or calcaneus with respect to the tibia. The similar results were expected, as the talus and calcaneus did not exhibit much motion relative to one another under passive loading. Quantifying moment arm as the distance between the Achilles tendon and center of the talar body (Csapo et al., 2010) was the second best predictor of geometric/MR (R^2^ = 0.50) and had an RMS error of less than 6 mm. External measurements of the malleoli and Achilles tendon measurement were positively correlated (R^2^ = 0.32) and within 6.5 mm of geometric/MR measurements on average. The length of the calcaneal tubercle (Raichlen et al., 2011) also positively correlated with geometric/MR, but over approximated geometric/MR measurements by 10 mm.

Tendon excursion did not correlate with geometric/MR measures of Achilles tendon moment arm, with the exception of passive tendon excursion/TE, which had a moderate correlation (R^2^ = 0.21) that approached statistical significance (p = 0.052). Attempting to control for tendon tension through submaximal contractions did not successfully improve agreement between tendon excursion/US and geometric/MR. Employing submaximal contractions appeared to increase the amount of variability in the measurements and for this reason may be counterindicated. Moment arms measured with tendon excursion/US measured during MVIC also did not correlate with those from geometric/MR but nonetheless may be linked to function (Lee and Piazza, 2009, 2012). While this measure may not reproduce moment arm measurements similar to geometric/MR it does quantify the amount of tendon travel during a maximal contraction, which is a surrogate measure of muscle shortening and tendon stiffness - two factors that are critical for locomotor function (Arampatzis et al., 2007; Kumagai et al., 2000).

### Alternative Bland-Altman plot

The traditional implementation of Bland-Altman plots is to demonstrate the agreement between two measurements: one accepted standard and one candidate replacement. While this was provided in the main body of the manuscript (Figure 3), the variance between the geometric/MR and tendon excursion/US were so great that we decided to plot the difference between these measures (Y-axis) against the geometric/MR (X-axis), which we consider to be the ‘gold standard’ of quantifying planar plantarflexor moment arms. Representing moment arm measurements using this modified Bland-Altman plot shows that there is no relationship between geometric/MR magnitude and difference between the two measurements.

**Appendix Figure 1.**
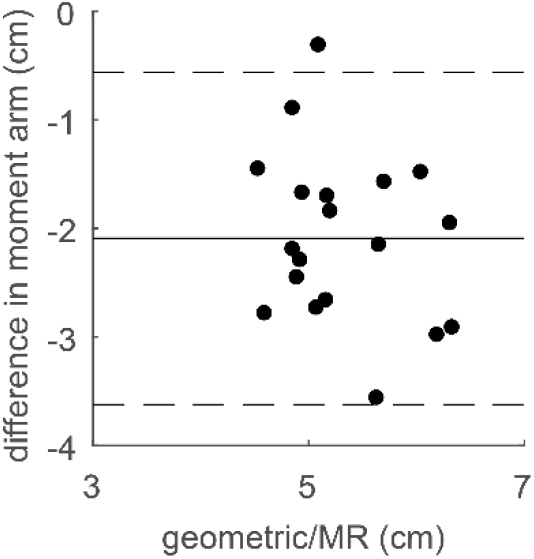
Tendon excursion/US measures consistently under approximated geometric/MR values of moment arm.

**Appendix Table 1.**

| Measurement | Type | Reference | R<sup>2</sup> (p-value) | RMS error (mm) |
| --- | --- | --- | --- | --- |
| Calcaneus - CoR | Geometric/MR | -- | 0.90 (<0.001) | 1.9 |
| Talus – circle fit | Geometric/MR | Csapo et al., 2010 | 0.50 (<0.001) | 5.7 |
| Malleolus distance | Geometric/External Measurement | Scholz et al., 2008 | 0.32 (0.012) | 6.5 |
| Tubercle length | Geometric/MR | Raichlen et al., 2011 | 0.23 (0.040) | 10.2 |
| TE – passive | Tendon Excursion/US | Waugh et al., 2011 | 0.21 (0.052) | 22.3 |
| TE – 30% MVIC | Tendon Excursion/US | Olszewski et al., 2015 | 0.09 (0.218) | 16.3 |
| TE – 60% MVIC | Tendon Excursion/US | -- | < 0.01 (0.930) | 18.0 |
| TE – 100% MVIC | Tendon Excursion/US | Lee and Piazza, 2009 | 0.06 (0.324) | 27.3 |

